# Drug synergy of combinatory treatment with remdesivir and the repurposed drugs fluoxetine and itraconazole effectively impairs SARS-CoV-2 infection in vitro

**DOI:** 10.1101/2020.10.16.342410

**Authors:** Sebastian Schloer, Linda Brunotte, Angeles Mecate-Zambrano, Shuyu Zheng, Jing Tang, Stephan Ludwig, Ursula Rescher

**Affiliations:** Institute of Medical Biochemistry, Center for Molecular Biology of Inflammation, and “Cells in Motion” Interfaculty Centre, University of Muenster, Von-Esmarch-Str. 56, D-48149, Muenster, Germany; Institute of Molecular Virology, Center for Molecular Biology of Inflammation, and “Cells in Motion” Interfaculty Centre, University of Muenster, Von-Esmarch-Str. 56, D-48149, Muenster, Germany; Research Program in Systems Oncology, Faculty of Medicine, University of Helsinki, Haartmaninkatu 8, 00029, Helsinki, Finland

## Abstract

The SARS-COV-2 pandemic and the global spread of coronavirus disease 2019 (COVID-19) urgently calls for efficient and safe antiviral treatment strategies. A straightforward approach to speed up drug development at lower costs is drug repurposing. Here we investigated the therapeutic potential of targeting the host- SARS-CoV-2 interface via repurposing of clinically licensed drugs and evaluated their use in combinatory treatments with virus- and host-directed drugs. We tested the antiviral potential of repurposing the antifungal itraconazole and the antidepressant fluoxetine on the production of infectious SARS-CoV-2 particles in the polarized Calu-3 cell culture model and evaluated the added benefit of a combinatory use of these host-directed drugs with remdesivir, an inhibitor of viral RNA polymerase. Drug treatments were well-tolerated and potent impaired viral replication was observed with all drug treatments. Importantly, both itraconazole-remdesivir and fluoxetine-remdesivir combinations inhibited the production of infectious SARS-CoV-2 particles > 90% and displayed synergistic effects in commonly used reference models for drug interaction. Itraconazole-Remdesivir and Fluoxetine-Remdesivir combinations are promising therapeutic options to control SARS-CoV-2 infection and severe progression of COVID-19.

## 1. INTRODUCTION

The zoonotic coronavirus SARS-CoV-2 and the resulting COVID-19 pandemic impressively show the global threat potential of a newly emerging pathogen. More than one million people have died so far from the current outbreak, and the proportion of infected people was estimated to reach more than 10% of the global population, with still unknown fatality rates (Baud et al., 2020; Rajgor et al., 2020; Wu et al., 2020). Because of the pressing burden on national health systems and economic losses, safe and efficient treatment strategies are urgently required. Developing a vaccine is a high priority. However, the rigorous testing and extensive clinical trials are time-consuming processes. While several trials are already ongoing, there are no vaccines available yet, and the required widespread access remains a challenging future task. Thus, approaches other than immunization might offer useful additional options for the management and control of SARS-CoV-2 infection and the treatment of COVID-19 (Fierabracci et al., 2020). A possibility to speed up the availability of drugs for the treatment of novel infections is the use of drugs that are already in clinical use for unrelated diseases via so-called “drug repurposing”. This approach represents a promising strategy to identify antiviral drugs with faster clinical implementation and lower development costs, considerations that are especially important in the global COVID-19 pandemic (Pushpakom et al., 2018). In addition to drugs that directly target the virus, host cell components that are vitally important in the viral life cycle are explored as promising starting points for therapeutic intervention (“host cell-directed therapy”) (Schwegmann and Brombacher, 2008; Ianevski et al., 2020; Zumla et al., 2020).

Although proteolytic cleavage of the SARS-CoV-2 spike protein surface protein by the host cell transmembrane protease serine 2 (TMPRSS2) enables SARS-CoV-2 to directly fuse with the plasma membrane, endocytosed SARS-CoV-2 particles use endosome-residing proteases for fusion within endosomes (Tang et al., 2020). Both pathways contribute to the SARS-CoV-2 infection process, and the preferential use of the actual fusion pathway might critically depend on the presence of plasma membrane proteases (Hoffmann et al., 2020). Our earlier research on influenza virus infection identified the late endosomal cholesterol balance as a critical factor in the influenza infection success and established this viral entry point as a possible pharmacological target. Elevated cholesterol levels inhibit the fusion of the influenza lipid hull with the endosomal membranes and thus inhibit the efficient transfer of the viral genome into the host cytosol (Musiol et al., 2013; Kühnl et al., 2018). We found that the clinically licensed antifungal itraconazole, a triazole derivative that blocks the fungal ergosterol pathway, has antiviral properties against a range of viruses and is effective against IAV infections in a preclinical mouse model (Schloer et al., 2019, 2020b). The additional therapeutic function is most likely based on direct inhibition of the endosomal cholesterol transporter Niemann-Pick Type C1 (NPC1) and the subsequent cholesterol storage ((Trinh et al., 2017; Schloer et al., 2019)).

The late endosome is an entry site for many zoonotically transmitted viruses, in particular for enveloped viruses including SARS-CoV-2 (Tang et al., 2020). Because of the functional similarities in transmitting the viral genome into the host cell, the same endosomal components might serve as pharmacological targets for a broad host-directed antiviral strategy against such viruses. Continuing our work on the endosomal host-virus interface, we explored whether a similar repurposing strategy could be used to impair SARS-CoV-2 entry and infection. Therefore, we assessed clinically licensed drugs that also affect endolysosomal lipid storage and cholesterol build-up for their antiviral potential. Here we report that itraconazole treatment potently inhibited the production of SARS-CoV-2 infectious particles. Together with our recently published work on the antiviral potential of the widely used serotonin inhibitor fluoxetine, which also negatively affects endosomal cholesterol release (Kornhuber et al., 2010; Schloer et al., 2020a) on SARS-CoV-2 infection, the results presented in this study strongly argue for the endolysosomal host-SARs-CoV-2 interface as a druggable target. However, host-directed drugs will rather suppress infection than completely eradicate the pathogen. The resulting demand for high drug doses and early and prolonged treatment is often associated with poor patient compliance. While drugs directly acting on virus structures are much more likely to completely eliminate the pathogens in shorter treatment time, emerging viral resistance to these antivirals is a major concern, as observed with the influenza neuraminidase inhibitor oseltamivir (Kim et al., 2013). Thus, combination therapies using virus- and host-directed drugs are considered to overcome these shortcomings. The nucleoside analog remdesivir which was originally developed against Ebola (Warren et al., 2016), has antiviral properties with a broad spectrum of activity against a number of RNA viruses and has already been shown to be effective against SARS and Mers-CoV in animal experiments (Sheahan et al., 2017, 2020; Agostini et al., 2018; Pruijssers et al., 2020). When remedesivir is incorporated into the viral RNA, the synthesis is prematurely terminated, and viral replication is inhibited (Gordon et al., 2020). Therefore, we explored combined treatments with remdesivir and the repurposed drugs itraconazole and fluoxetine. Both drug combinations showed stronger antiviral activities against influenza viruses compared to remdesivir monotherapy (Schloer et al., 2020b). Importantly, the antiviral effects of the combined treatments deviated from the expected effects, and pharmacodynamic evaluation via commonly used reference models to study drug interaction revealed synergistic interaction.

## 2. MATERIALS AND METHODS

### 2.1 Cells and SARS-CoV-2 isolate

The human bronchial epithelial cell lines Calu-3 and the Vero E6 cells were cultivated in Dulbecco’s modified Eagle’s medium (DMEM) with 10% standardized fetal bovine serum (FBS Superior; Merck), 2 mM L-glutamine, 100 U/mL penicillin, 0.1 mg/mL streptomycin, and 1% non-essential amino acids (Merck) in a humidified incubator at 5% CO_2_ and 37 °C. Calu-3 monolayers were polarized and cultured as described (Schloer et al., 2020b). Itraconazole (2 mg/mL, Sigma), fluoxetine (5 mM, Sigma) and remdesivir (10 mM, Hycultec) were solubilized in DMSO. The SARS-CoV-2 isolate hCoV-19/Germany/FI1103201/2020 (EPI-ISL_463008, mutation D614G in spike protein) was amplified on Vero E6 cells and used for the infection assays.

### 2.2 Cytotoxicity assay

Calu-3 cells were cultured at the indicated concentrations with either the solvent DMSO, itraconazole or with the combinations of itraconazole/ remdesivir (ItraRem) or fluoxetine/ remdesivir (FluoRem) for 48 h. Staurosporine solution (1 μM) served as a positive control for cytotoxic effects. After 48 hours of treatment, cell viability was evaluated by adding MTT 3-(4,5-dimethylthiazol-2-yl)-2,5-diphenyltetrazolium bromide (Sigma) to the cells for 4 h and OD_562_ measurements according to the manufacturer’s protocols (Sigma).

### 2.3 Inoculation of cells and drug treatment

For infection, polarized Calu-3 and Vero cells were washed with PBS and inoculated at a multiplicity of infection (MOI) of 0.1 (Calu-3) or 0.01 (Vero E6) of virus diluted in infection-PBS (containing 0.2% BSA, 1% CaCl_2_, 1% MgCl_2_, 100 U/mL penicillin and 0.1 mg/mL streptomycin) at 37°C for 1 hours. Subsequently, cells were washed with PBS and further cultured in infection-DMEM (serum-free DMEM containing 0.2% BSA, 1 mM MgCl_2_, 0.9 mM CaCl_2_, 100 U/mL penicillin, and 0.1 mg/mL streptomycin) at 5% CO_2_ and 37 °C. Cells were treated with the indicated remdesivir, itraconazole or fluoxetine concentration 2 hours post-infection (hpi) for the entire 48 hours infection period. After 48 hpi, apical culture supernatants were collected and immediately frozen at −80 °C until titration via plaque assay.

### 2.4 Plaque assay

To determine the number of infectious particles in the supernatant of treated cells, a standard plaque assay was performed. Briefly, Vero E6 cells grown to a monolayer in six-well dishes were washed with PBS and infected with serial dilutions of the respective supernatants in infection-PBS for 1 hour at 37 °C. The inoculum was replaced with 2x MEM (MEM containing 0.2% BSA, 2 mM L-glutamine 1 M HEPES, pH 7.2, 7.5% NaHCO3, 100 U/mL penicillin, 0.1 mg/mL streptomycin, and 0.4% Oxoid agar) and incubated at 37°C. Virus plaques were visualized by staining with neutral red, and virus titers were calculated as plaque-forming units (PFU) per mL.

### 2.5 Data and statistical analysis

A priori power analysis using G*Power 3.1 (Faul et al., 2007)) was performed to determine the sample sizes required to detect > 90% reduction in virus titers at powers > 0.8. Data were analyzed using the software GraphPad Prism version 8.00 (GraphPad). No outliers were detected.

Infectious viral titers are presented as means ± SEM of five measurements per experiment per condition. For dose-response curves, drug concentrations were log-transformed and virus titers were expressed as percentages of the mean virus titer in control cells (treated with the solvent DMSO) and data were analyzed by curve fitting using a 4 parameter logistic model. Drug combinatory effects were analyzed by using SynergyFinder, an open-source free stand-alone web application for the analysis of drug combination data (Ianevski et al., 2017). Synergy was evaluated based on the Zero Interaction Potency (ZIP), Bliss independence, and Highest single agent (HSA) reference models (He et al., 2018). We further analyzed the overall drug combination sensitivity score (CSS) by using the CSS method (Malyutina et al., 2019).

## 3. RESULTS

### 3.1. The clinically licensed antifungal drug itraconazole efficiently blocks the production of SARS-CoV-2 infectious particles

Based on the successful use of repurposing itraconazole for the treatment of influenza virus infection reported in our earlier studies (Schloer et al., 2019, 2020b), we first established whether this clinically licensed drug also had an antiviral potential on the production of infectious SARS-CoV-2 particles. In line with our previous results (Schloer et al., 2020a), both Calu-3 and Vero E6 cell lines supported SARS-CoV-2 replication and produced high virus titers (Fig. 1). Inoculation with the clinical isolate hCoV-19/Germany/FI1103201/2020 at MOI 0.1 yielded 1.46× 10^6^ PFU in Calu-3 cells and 6.37× 10^6^ PFU at MOI 0.01 in Vero cells at 48 hpi. Treatment with itraconazole 2 hpi potently inhibited SARS-CoV-2 replication in a dose-dependent manner in both cell types (Fig. 1a). Fitting the experimental dose-response values to a nonlinear 4 parameter logistic model revealed half-maximal inhibitory (EC_50_) and 90% inhibitory concentrations (EC_90_) of 0.43 μM and 2.46 μM in Calu-3 cells, and even lower 50% and 90% inhibitory concentrations (0.39 μM and 0.87 μM) were determined for itraconazole antiviral activity in SARS-CoV-2 infected Vero cells (Fig. 1b). Of note, no detectable cytotoxicity was observed with these doses (Fig. S1a). The itraconazole cytotoxic concentration required to reduce cell viability by 50% (CC_50_) was determined at 25.56 μM, resulting in a selectivity index (SI, defined as the ratio of CC_50_ to EC_50_) of 25.56 μM which indicated an effective and safe antiviral window.

**FIGURE 1.**
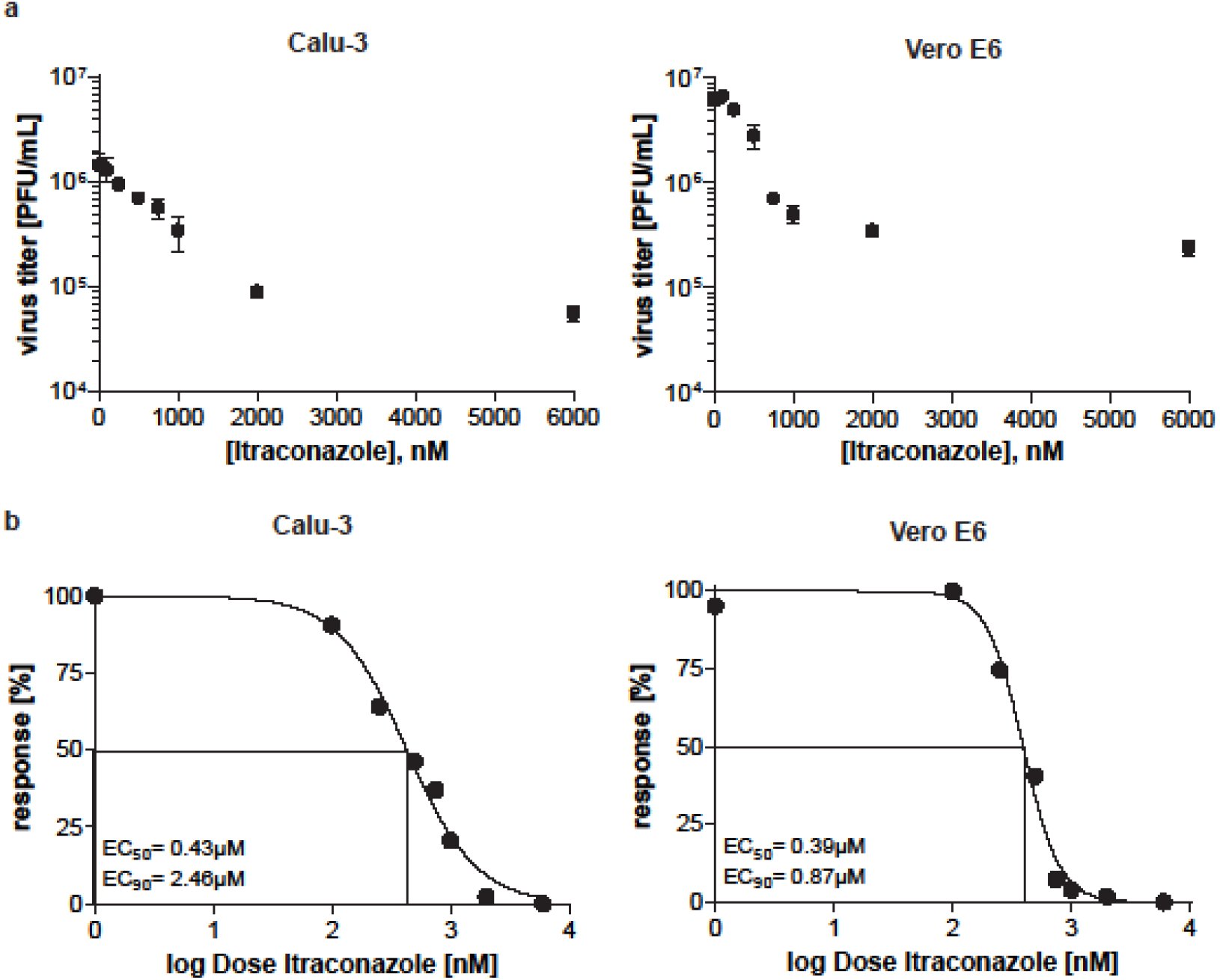
Analysis of itraconazole-mediated reduction of SARS-CoV-2 replication. Calu-3 and Vero E6 cells were infected with 0.1 MOI of SARS-CoV-2. At 2 hpi, cells were treated with itraconazole at the indicated concentrations. (a) Mean infectious viral titers□±□SEM, (b) Mean percent inhibition ± SEM of SARS-CoV-2 replication, with mean virus titer in control cells (treated with the solvent DMSO) set to 100%; n□=□5. LogEC_50_ and LogEC_90_ values were determined by fitting a non-linear regression model. (Calu-3: EC_50_ = 0.43 μM, EC_90_ = 2.46 μM; Vero E6: EC_50_ = 0.39 μM, EC_90_ = 0.87 μM).

### 3.2 Remdesivir inhibits SARS-CoV-2 replication in polarized Calu-3 cells

We addressed the question of whether a host-targeting drug could be used in combination with a virus-directed drug for more efficient suppression of SARS-CoV-2 replication. Thus, we first assessed the antiviral capacity of remdesivir, a nucleotide analog prodrug that inhibits SARS-CoV-2 viral RNA-dependent RNA polymerase (Gordon et al., 2020). The EC_50_ concentration was reached at 0.42 μM and EC_90_ at 1.08 μM in Calu-3 cells (Fig. S2), well in line with published data (Pruijssers et al., 2020). Next, we determined viral replication in cells that had been treated with combinatory therapy. Because we recently published the potential use of repurposing the antidepressant fluoxetine for treatment of SARS-CoV-2 infection (Schloer et al., 2020a), we also assessed the effects of a combined fluoxetine/remdesivir (FluoRem) treatment in addition to the itraconazole/remdesivir (ItraRem) combination (Fig. 2). Both drugs are clinically licensed and do not induce significant cytotoxicity over the whole concentration range ((Schloer et al., 2020a), and Fig. S1a). The combination treatments were also well-tolerated, and no cytotoxic effects were seen when cells were simultaneously treated with the drug pairs (Fig. S1b, c).

**FIGURE 2.**
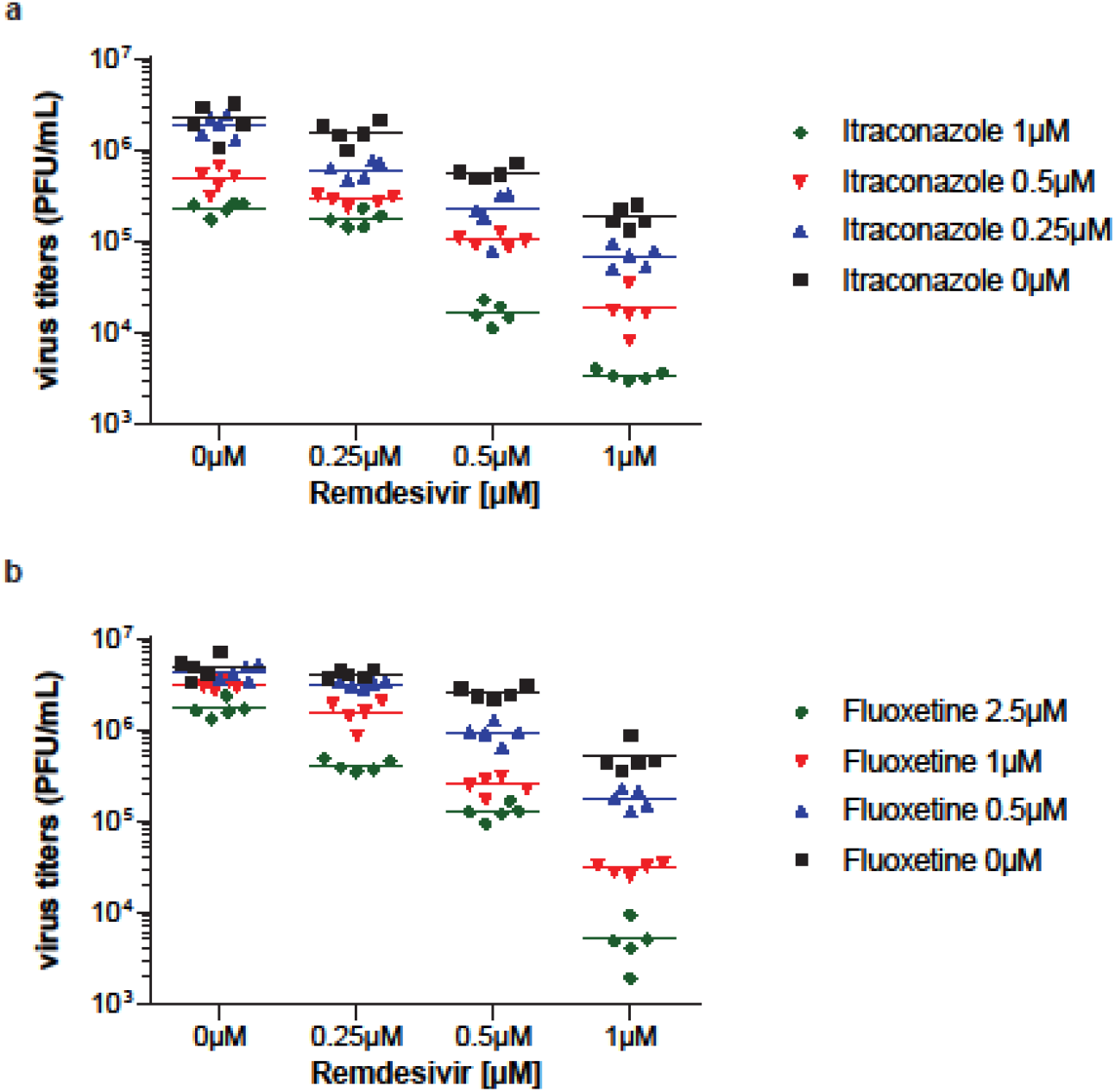
Antiviral activities of treatments. Infectious virus production in Calu-3 cells treated as indicated 2 hpi. Each symbol represents plaque-forming units (PFU) per mL detected in a single experimental sample, lines indicate means.

### 3.3 Combinatory treatments with the drug pairs itraconazole-remdesivir or fluoxetine-remdesivir show enhanced antiviral activity due to synergistic interaction

For all drugs, we chose those concentrations that were not sufficient to achieve a 90% reduction when individually applied (Fig 3a). For both ItraRem and FluoRem combinations, a potent reduction in virus titers was detected in all cases. Of note, several combinations yielded a reduction > 90% of the maximum virus titers produced in control cells (Fig. 3b).

**FIGURE 3.**
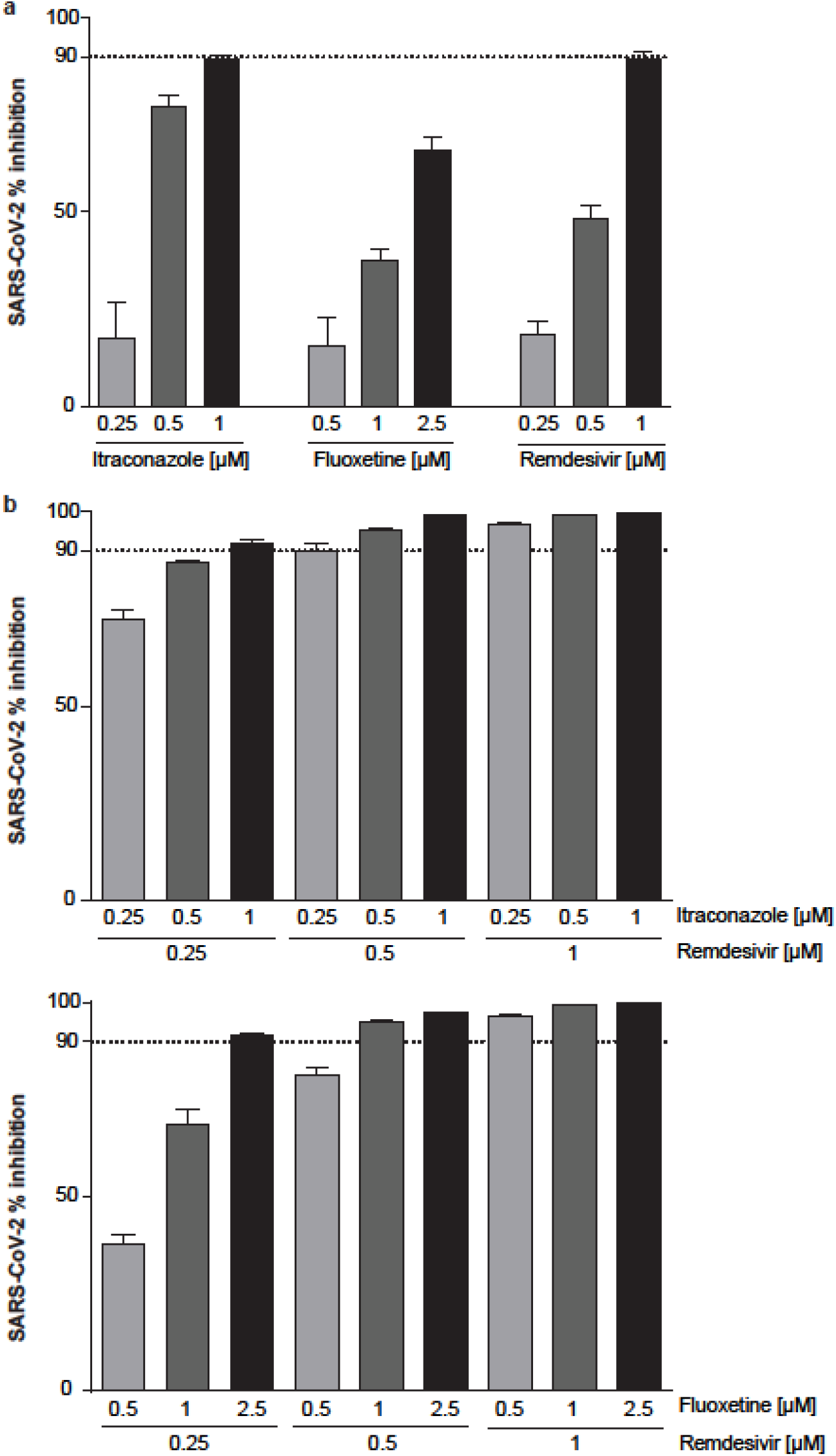
Antiviral activities of single and combination treatments. Calu-3 cells were treated with the indicated drug combinations for 48 h. (a) Single treatment, (b) combinatory treatment. Bars represent mean percent inhibition□±□SEM of infectious virus production, with mean virus titer produced in control cells (treated with the solvent DMSO) set to 100%; n□=□5. Dotted line, 90% reduction in viral titer.

We, next considered the pharmacological interactions of the respective drug pairs. Thus, we evaluated the drug interactions via Bliss independence, Highest single agent (HSA), and ZIP, three commonly used reference synergy models that differ in their basic assumptions of drug interaction they are based on. The results, presented in Figs. 4 and 5, consistently argued for synergistic action of remdesivir with itraconazole and fluoxetine, as indicated by the positive average synergy score across all models. Closer inspection of the drug interaction relationships and landscape visualizations revealed that for ItraRem, the highest synergy scores were calculated with the lower concentration ranges of both drugs (Fig. 4). The strong synergy led to an overall drug combination sensitivity score (CSS) of 89.64, resulting in >90% inhibition already at 500 nM of remdesivir and 250 nM of itraconazole. FluoRem combination treatment had a higher average synergy score, as well as a higher CSS score than ItraRem (92.82 vs 89.64), suggesting that this drug combination is more likely to show synergy. Importantly, for all models, the FluoRem combinations that met the ≥ 90% inhibition criterion were well within the high synergy area (Fig. 5).

**FIGURE 4.**
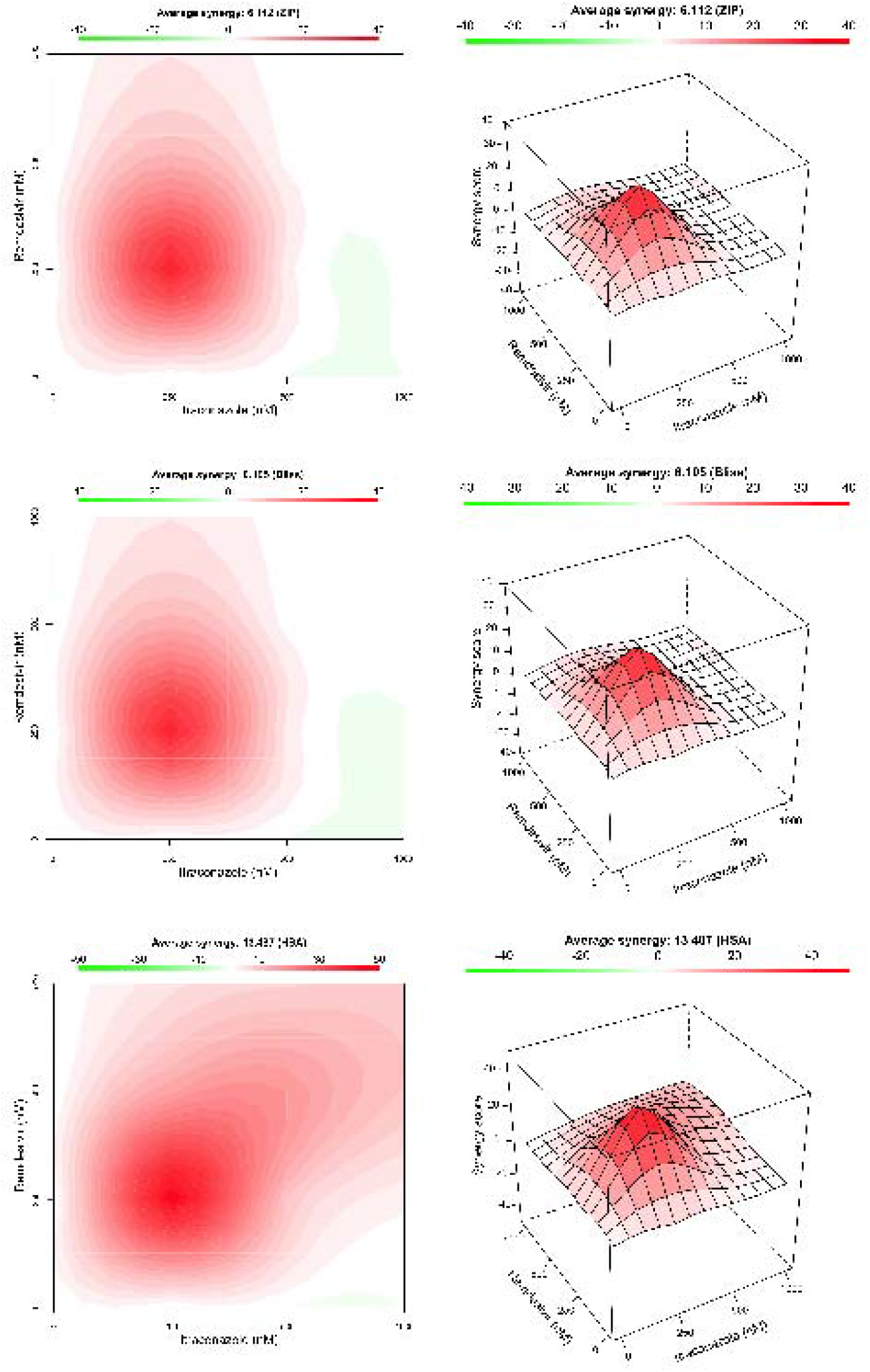
Evaluation of the pharmacological interactions of itraconazole and remdesivir (ItraRem). ZIP, Bliss independence, and Highest single agent (HSA) reference models were used to assess the interaction landscapes. Interaction surfaces are color-coded according to the synergy scores of the responses.

**FIGURE 5.**
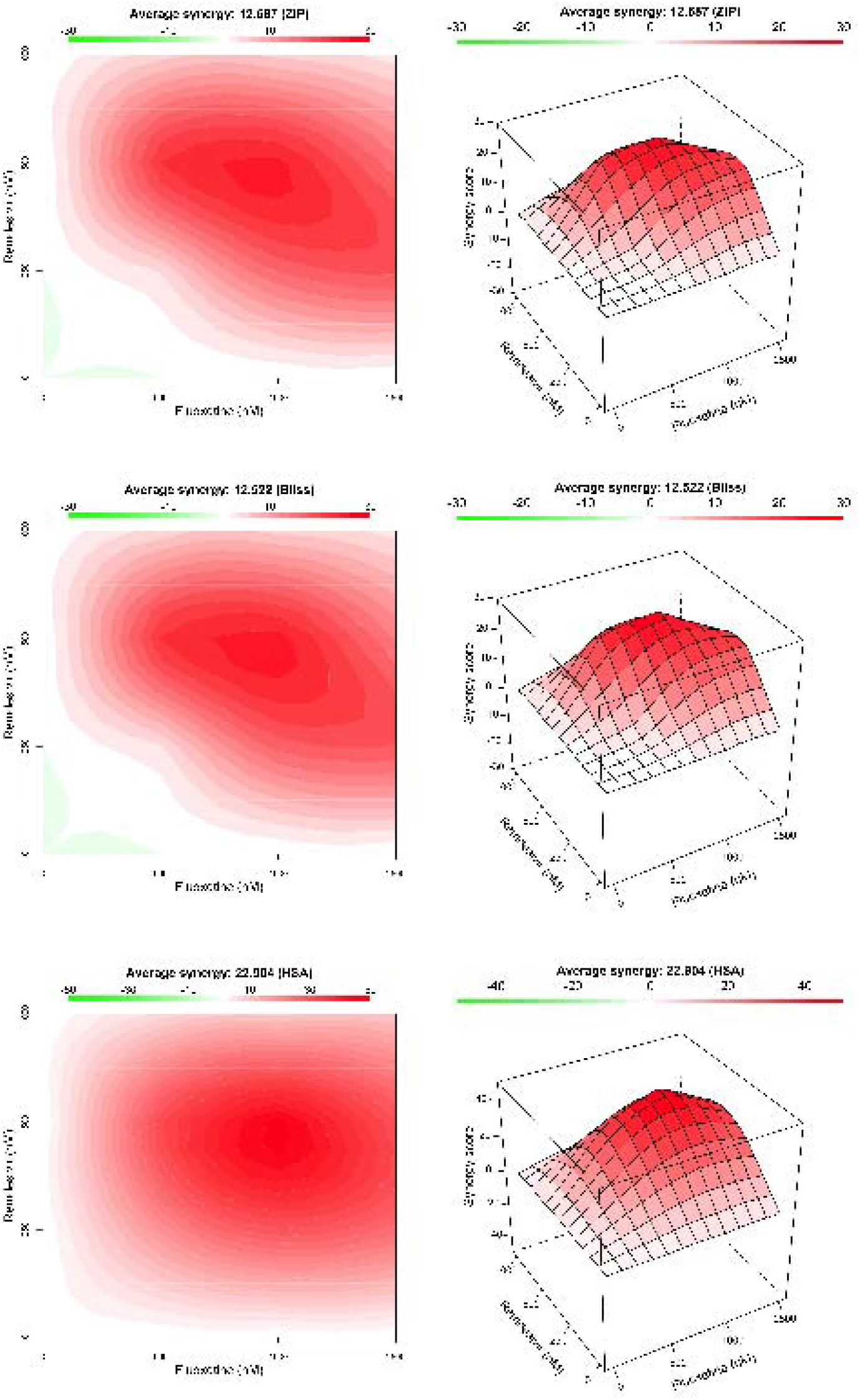
Evaluation of the pharmacological interactions of fluoxetine and remdesivir (FluoRem). ZIP, Bliss independence, and Highest single agent (HSA) reference models were used to assess the interaction landscapes. Interaction surfaces are color-coded according to the synergy scores of the responses.

## 4. DISCUSSION

The nucleoside analog remdesivir with broad-spectrum antiviral activity against a range of viruses including Ebola, Marburg, MERS, and SARS inhibits viral RNA polymerase and is also active against SARS-CoV-2 (Warren et al., 2016; Sheahan et al., 2017, 2020; Agostini et al., 2018; Brown et al., 2019). Indeed, remedesivir was the first new drug to receive an FDA emergency use authorization for the treatment of severe COVID-19 cases. However, a common concern about virus-directed antivirals is the development of drug resistance. Due to their error-prone replication mode, drug-resistant viruses are increasingly encountered (Strasfeld and Chou, 2010), as observed with the influenza neuraminidase inhibitor oseltamivir (Kim et al., 2013). Because profound changes would be required to allow viruses to replicate independently of otherwise essential host factors, the development of host-directed therapeutics is an emerging concept. However, host-directed drugs will rather cause impaired viral replication than complete eradication, thus demanding high drug concentrations and starting treatment start as early as possible and for extended durations, which is often associated with poor patient compliance. Because of these shortcomings, combinatory treatments with both virus- and host-directed drugs are explored for enhanced treatment success.

Enveloped viruses such as SARS-CoV-2 depend on the fusion of their lipid hull with the host membrane to get access to the cytosol. The SARS-CoV-2 spike protein, which protrudes from the virus surface, mediates initial binding to angiotensin-converting enzyme 2 (ACE2), which serves as the host cell surface receptor (Li et al., 2003; Lan et al., 2020; Ou et al., 2020; Zhou et al., 2020). To promote fusion with the host cell membrane, the spike protein needs to be primed by proteolytic cleavage, which can be mediated by several host proteases. Transmembrane protease serine 2 (TMPRSS2)-mediated cleavage leads to fusion with the plasma membrane, while endosome-residing proteases are utilized by endocytosed SARS-CoV-2 particles for fusion within endosomes. Since both routes have been reported to contribute to the SARS-CoV-2 infectivity (Hoffmann et al., 2020), the endosomal compartment is also a critical host/pathogen interface for SARS-CoV-2. Our previous studies strongly support the late endosomal cholesterol balance as a cellular target for antiviral intervention (Musiol et al., 2013; Kühnl et al., 2018; Schloer et al., 2019, 2020a, 2020b). Notably, endolysosomal lipid storage and cholesterol build-up could be induced via repurposing of drugs approved for unrelated applications. We found that the triazole derivative itraconazole, a clinically licensed antifungal drug (Organization, 2019) that directly inhibits the endosomal cholesterol transporter NPC1 (Trinh et al., 2017) has an antiviral potential on the endosomal fusion of enveloped viruses including influenza viruses (Schloer et al., 2019). The findings presented in this study add SARS-CoV-2 to the spectrum of itraconazole-sensitive enveloped viruses. Our results reveal a potent antiviral activity of itraconazole on the production of SARS-CoV-2 infectious particles, with EC50 values comparable to what we previously reported for itraconazole-mediated antiviral activity against IAV subtypes. The bioavailability after oral application of itraconazole is low because of the low water solubility of this compound (Grant Prentice and Glasmacher, 2005; Domínguez-Gil Hurlé et al., 2006). Only limited amounts are absorbed from the gastrointestinal tract after ionization, and levels depend on the individual gastric acidity (Shin et al., 2004; Domínguez-Gil Hurlé et al., 2006; Bae et al., 2011; Lestner and Hope, 2013; Allegra et al., 2017). However, our previous study revealed a beneficial antiviral activity in vivo (Schloer et al., 2019).

We recently discovered that the widely used antidepressant fluoxetine has strong SARS-CoV-2 antiviral activity (Schloer et al., 2020a). Together with the findings presented here on the inhibitory function of itraconazole treatment on SARS-CoV-2 replication, these results suggest that both drugs are promising candidates for repurposing as a host-directed drug for SARS-CoV-2 infection treatment. Both drugs most likely interfere with the proper endosomal cholesterol levels. Whereas itraconazole und posaconazole both directly inhibit the endosomal cholesterol transporter NPC1 (Trinh et al., 2017), fluoxetine functionally blocks the endolysosome-residing enzyme sphingomyelin phosphodiesterase (“acid sphingomyelinase”, ASM), which in turn causes sphingomyelin accumulation and negatively affects cholesterol release from this compartment (Kornhuber et al., 2010).

## 5. CONCLUSION

While remdesivir and the host-directed drugs itraconazole or fluoxetine target independent pathways, we found that drug combinations together with remdesivir (ItraRem and FluoRem) showed stronger antiviral activities against SARS-CoV-2 than the remdesivir monotherapy. Moreover, the overall therapeutic effect of the combinations was larger than the expected sum of the independent drug effects. Our analysis on the antiviral activity of combinatory drug combinations via commonly used interaction models argue for an enhanced efficacy that is based on synergistic drug interaction and suggests promising novel options for SARS-CoV-2 treatment.

## ACKNOWLEDGEMENTS

We thank Jonathan Hentrey for help with the plaque assays.

## AUTHOR CONTRIBUTIONS

Conceptualization and methodology, S.S., U.R.; validation, formal analysis, investigation, data curation, S.S., J.T., U.R.; resources, J.T., S.L.; U.R.; writing—original draft preparation, U.R.; writing—review and editing, S.S., S.L., U.R.; visualization, S.S.; supervision, U.R., S.L., project administration, U.R.; funding acquisition, U.R. All authors have read and agreed to the published version of the manuscript.

## SUPPORTING INFORMATION

**SUPPORTING FIGURE 1.**
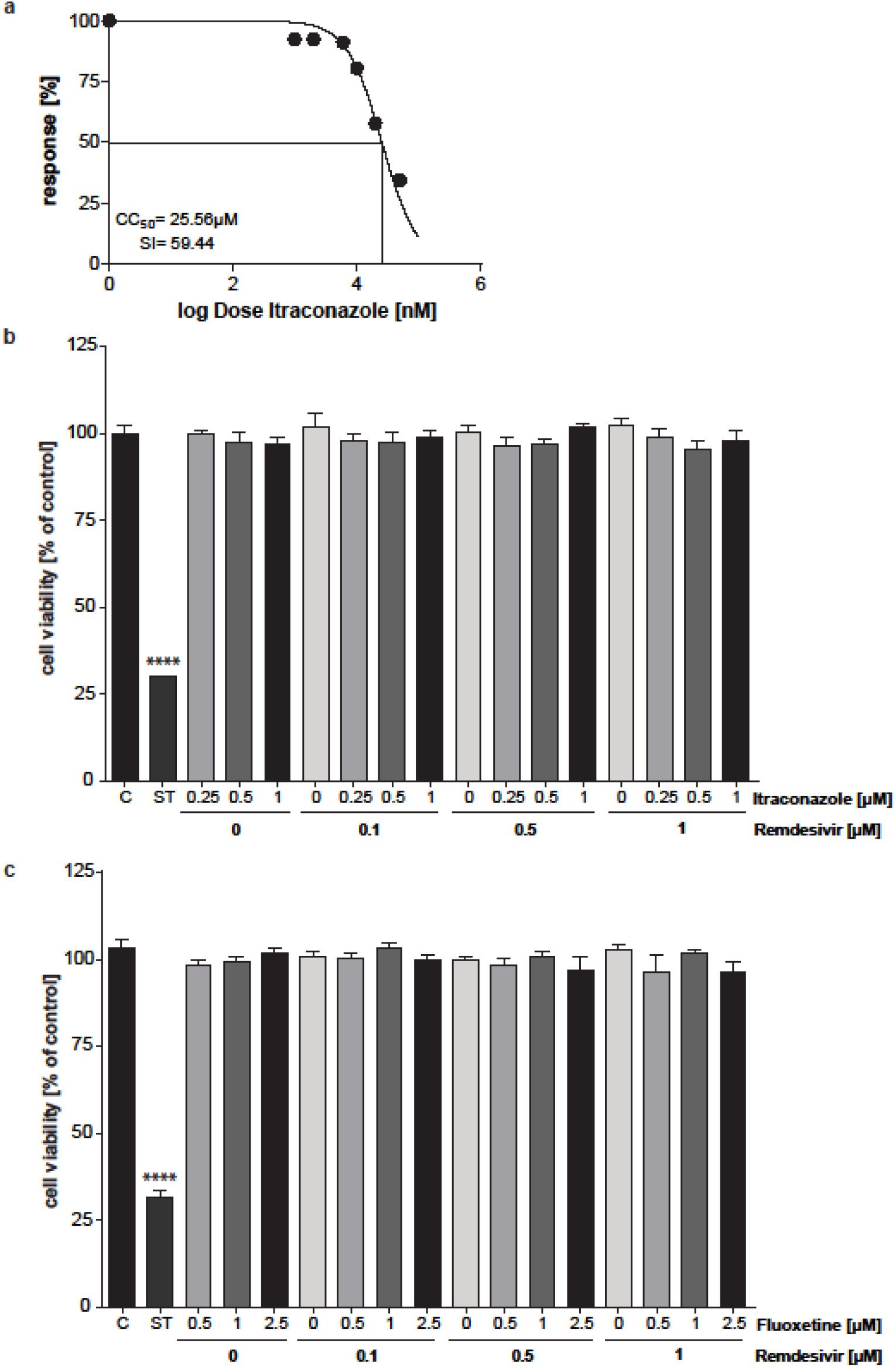
Analysis of the cytotoxicity of treatments. (a) Itraconazole, (b) ItraRem, and (c) FluoRem combinatory treatment. Calu-3 cells were treated with the indicated drug combinations for 48 h. Bars display mean percentages of viable cells□±□SEM, with mean viability in solvent-treated control cells (C) set to 100%. Staurosporine (ST)-induced cytotoxicity served as a positive control. n□=□5, one-way ANOVA followed by Dunnett’s multiple comparison test, ****p□<□0.0001.

**SUPPORTING FIGURE 2.**
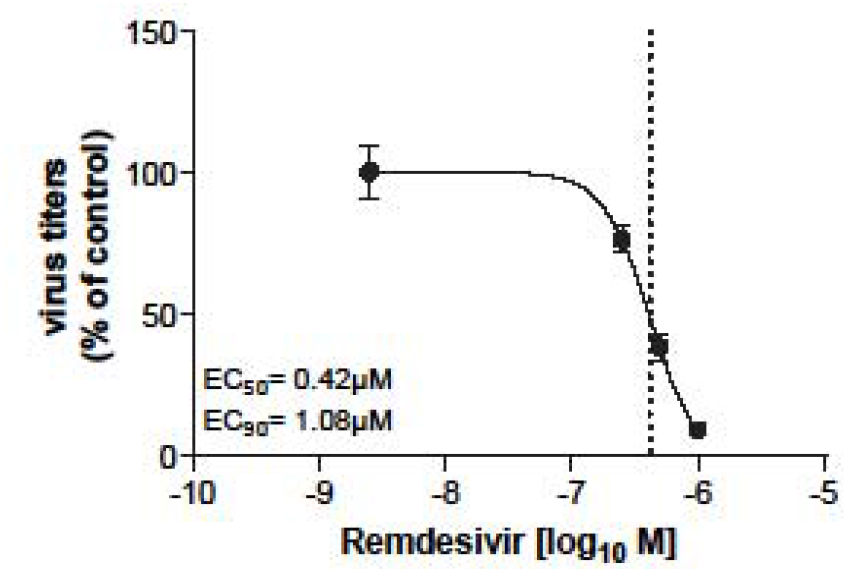
Analysis Dose-Response curve of remdesivir treatments in Calu-3 cells. Calu-3 cells were infected with 0.1 MOI of SARS-CoV-2 for 1 h and treated with the indicated drug combinations for 48 h. Mean percent inhibition ± SEM of SARS-CoV-2 replication, with mean virus titer in control cells (treated with the solvent DMSO) set to 100%; n□=□5. LogEC_50_ and LogEC_90_ values were determined by fitting a non-linear regression model. (Calu-3: EC50 = 0.42 μM, EC90 = 1.08 μM).

